# Mitochondrial fostering: the mitochondrial genome may play a role in plant orphan gene evolution

**DOI:** 10.1101/2020.01.02.884874

**Authors:** Seth O’Conner, Ling Li

## Abstract

Plant mitochondrial genomes exhibit odd evolutionary patterns. They have a high rearrangement but low mutation rate, and a large size. Based on massive mitochondrial DNA transfers to the nucleus as well as the mitochondrial unique evolutionary traits, we propose a “Mitochondrial Fostering” theory where the organelle genome plays an integral role in the arrival and development of orphan genes (genes with no homologues in other lineages). Two approaches were used to test this theory: 1) bioinformatic analysis of nuclear mitochondrial DNA (Numts: mitochondrial originating DNA that has migrated to the nucleus) at the genome level, and 2) bioinformatic analysis of particular orphan sequences present in both the mitochondrial genome and the nuclear genome of *Arabidopsis thaliana*. One study example is given about one orphan sequence that codes for two unique orphan genes: one in the mitochondrial genome and another one in the nuclear genome. DNA alignments show regions of this *A. thaliana* orphan sequence exist scattered throughout other land plant mitochondrial genomes. This is consistent with the high recombination rates of mitochondrial genomes in land plants. This may also enable the creation of novel coding sequences within the orphan loci, which can then be transferred to the nuclear genome and become exposed to new evolutionary pressures. Our study also reveals a high correlation between mitochondrial DNA rate transferred to the nuclear genome and number of orphan genes in land plants. All the data suggests the mitochondrial genome may play a role in nuclear orphan gene evolution in land plants.

## Introduction

Our depth of understanding organism genomes has progressed with the advancement of sequencing technology. Throughout all domains of life, there is thought to be some level of species-specific genes called orphan genes (Tautz & Domazet-Loso, 2011). Around 5-15% of an organism’s genes are estimated to be orphans (Arendsee et al., 2014). The function of orphan genes can range from negligible to necessary; mutants of some orphan genes in Drosophila cause lethality (Reinhardt et al., 2013). Orphan genes may also explain some species-specific functionality—Prod1, a salamander specific gene, aids in limb regeneration of salamanders, the only tetrapod with this ability (Kumar et al., 2015). In *Arabidopsis thaliana*, Qua Quine Starch (QQS) - the species-specific orphan gene and its network have been shown to be involved in regulation of carbon and nitrogen allocation (Li et al., 2009; Li & Wurtele, 2015; Li et al., 2015; Jones et al., 2016), and several orphans have been implicated in stress response (Luhua et al., 2013; Qi et al., 2018; Graham et al., 2004). Since the discovery of orphan genes in the yeast genome in 1996 (Dujon et al., 1996), researchers have made strides to explain the origin of these novel genes. However, the orphan gene origin story is still under debate today (Rodelsperger, 2018).

It was previously thought the main mechanism involved in developing novel genes was duplication and subsequent sequence diversification (Tautz & Domazet-Loso, 2011). However, as put forth by Tautz and Domazet-Loso (2011), there seems to be more and more evidence for *de novo* evolution of genes out of previously non-coding sequences. Although there are several theories for how *de novo* genes evolve (Bennetzin, 2005), it is not clear how a previously non-coding sequence can develop into a gene, let alone a functional gene. Many speculate *nuclear* non-coding DNA to be the substrate through which gene evolution occurs (Bennetzin, 2005). However, it is important to consider all “nucleotide banks” (*i.e.*, organelle genomes) when trying to pinpoint the source of new genes. Noutsos et al., 2007 determined a possible role for Numt (mitochondrial originating DNA that has migrated to the nucleus) DNA in exon evolution. This work showed a small sampling of Numt DNA had been functionally integrated into exons of nuclear genes. They proposed there were likely more Numt insertions into exon regions that were now unrecognizable due to sequence divergence. While some work has been done to show that Numts may play a role in remodeling nuclear genes (Noutsos et al., 2007), there is currently little research into mitochondrial genomes and their impact on *de novo* orphan gene evolution.

The classical evolutionary theory of endosymbiosis states that modern eukaryotic cells are the product of a symbiotic event, wherein a proteobacterium was engulfed by a larger cell which eventually became the eukaryotic cell (Timmis et al., 2004). These proteobacteria became the mitochondria, which contains its own genome. Since the endosymbiotic event, the mitochondrial genome has begun to transfer its genes into the nuclear genome (Berg & Kurland, 2000). To put into perspective just how massive this gene transfer event has been, it is thought that there were once 1,600 proteins coded within the proto-mitochondrial cell, but now in modern mitochondria, there are typically 67 proteins or less coded from the organelle genome (Berg & Kurland, 2000). This process of mitochondrial genetic material transfer has been ongoing in the form of Numts (Timmis et al., 2004).

Currently, mitochondrial genomes in plants are known for odd evolutionary patterns. They are large and rearranged at a high rate while having a low mutation rate (Christensen, 2013). They also contain long intergenic regions—up to 98% (Kubo & Mikami, 2006; Skippington et al., 2015; Sloan et al., 2012). It has been recently shown that large changes between mitochondrial genomes in various Arabidopsis ecotypes are due to high recombinatorial activity controlled by *MSH1,* a nuclear gene responsible for suppressing recombination of mitochondrial DNA repeats (Arrieta-Montiel et al. 2009). The novelty of mitochondrial sequences in plant species is possibly due to large repeats and rearrangements, sequence influx from the nuclear and chloroplast genomes, and horizontal sequence transfer from other plant mitochondria (Wynn & Christensen, 2018; Rice et al., 2013; Sanchez-Puerta et al., 2008; Xi et al., 2013). Based on large mitochondrial nucleotide transfers to the nucleus of current species (Hazkani-Covo et al., 2010; Zhao et al., 2018) as well as the mitochondrial unique evolutionary traits, we analyzed Numt and orphan gene data for six plant species and six animal species as well as a particular young gene present in both the mitochondrial genome and nuclear genome of *A. thaliana*. We propose a “Mitochondrial Fostering” theory where the plant mitochondrial genome may play an integral role in the arrival and development of *de novo* orphan genes. This process appears to be a continuation of the gene transfer events that began to occur after endosymbiosis (Berg & Kurland, 2000). Two approaches were used to test this theory: 1) Bioinformatic analysis of Numts at the genome level and 2) bioinformatic analysis of particular mitochondrial orphan sequences that are present in both the mitochondrial genome and the nuclear genome of *A. thaliana*—and one that codes for two different genes: AtMg01180 in the mitochondrial genome and At2g07667 in the nuclear genome on chromosome two.

## Results

### *Arabidopsis thaliana*, *Glycine max*, and *Zea mays* contain a large proportion of orphan genes in mitochondrial genome

To determine which cellular genomes have the highest orphan gene potential, we analyzed the number of orphan genes in mitochondrial and whole genomes (*i.e.*, all nuclear, mitochondrial and chloroplast genomes) in three plant species. *G. max* has the largest number of total genes in the genome, while *Z. mays* has the largest number of mitochondrial genes, mitochondrial orphan genes, and total orphan genes (from all genomes) (Table 1). *Z. mays* also has the highest proportion of orphan genes for the whole genome and the largest proportion of orphan genes in the mitochondrial genome (Fig. 1). All three species have a much higher proportion of orphan genes in the mitochondrial genome compared to the proportion in the whole genome (Fig. 1). To understand if the phenomenon is prevalent in all organelle genomes, orphans were determined for the chloroplast genomes as well. Similar proportions are present in both the whole genome and the chloroplast genome for *A. thaliana* and *G. max* while *Z. mays* has a higher proportion in the whole genome compared to the chloroplast genome (Fig. 1). As there is not a large proportion of orphan genes in the chloroplast genomes, the mitochondrial genome was focused on for this study.

**Figure 1.**
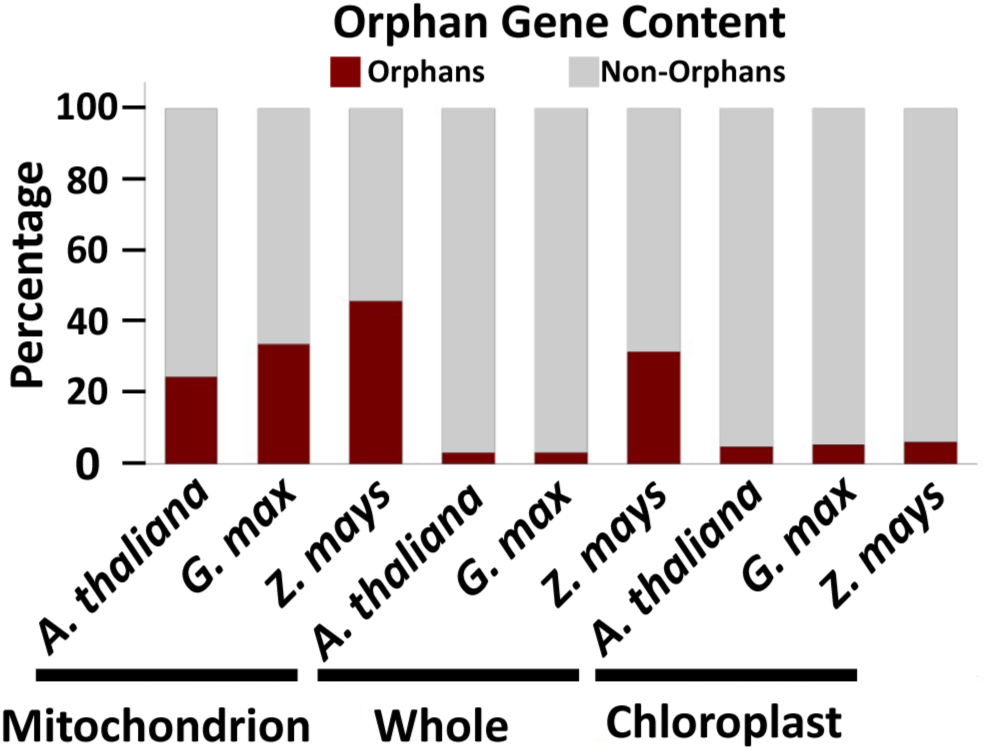
Orphan gene content in three well characterized land plants. Three plant genomes show higher orphan gene content in mitochondrial genome compared to whole genome and chloroplast genome

**Table 1.** Comparison of the three cellular genomes in A. thaliana, G. max and Z. mays. All three species have more mitochondrial orphan genes compared to chloroplast orphans.

|  | **Whole**<br>**genome**<br>**orphan** | **Mitochondrion**<br>**orphans** | **Chloroplast**<br>**orphans** | **Total**<br>**genes** | **Total**<br>**mitochondrial**<br>**genes** | **Total**<br>**chloroplast**<br>**genes** |
| --- | --- | --- | --- | --- | --- | --- |
| *A.*<br>*thaliana* | 1,169 | 30 | 4 | 35,387 | 122 | 80 |
| *G. max* | 2,410 | 24 | 5 | 73,320 | 71 | 79 |
| *Z. mays* | 5,245 | 63 | 5 | 63,540 | 137 | 91 |

### Orphan genes are predicted to preferentially locate to mitochondria in *A. thaliana*

As nuclear mitochondrial genes must gain a mitochondrial targeting peptide to localize back to the mitochondria (Berg & Kurland, 2000), we determined the proportion of orphan genes with predicted mitochondrial targeting peptides to determine if orphan genes preferentially code for mitochondrial targeting peptides, possibly signifying a mitochondrial origin. Subcellular localization data for *A. thaliana* was obtained from TAIR (arabidopsis.org) and cross-referenced with our *A. thaliana* orphan gene list (Arendsee et al., 2014). The largest portion (47%) of *A. thaliana* orphans had predicted localization to mitochondria, which was even larger than unknown localization (37.5%) (Fig. 2A). To verify this analysis, the component GO terms were downloaded for 852 orphan genes that have the component GO terms. Again, mitochondrion had the largest number of hits (419) with unknown in second (209 hits)—percentages of 47 and 23.4 respectively (Fig. 2B).

**Figure 2.**
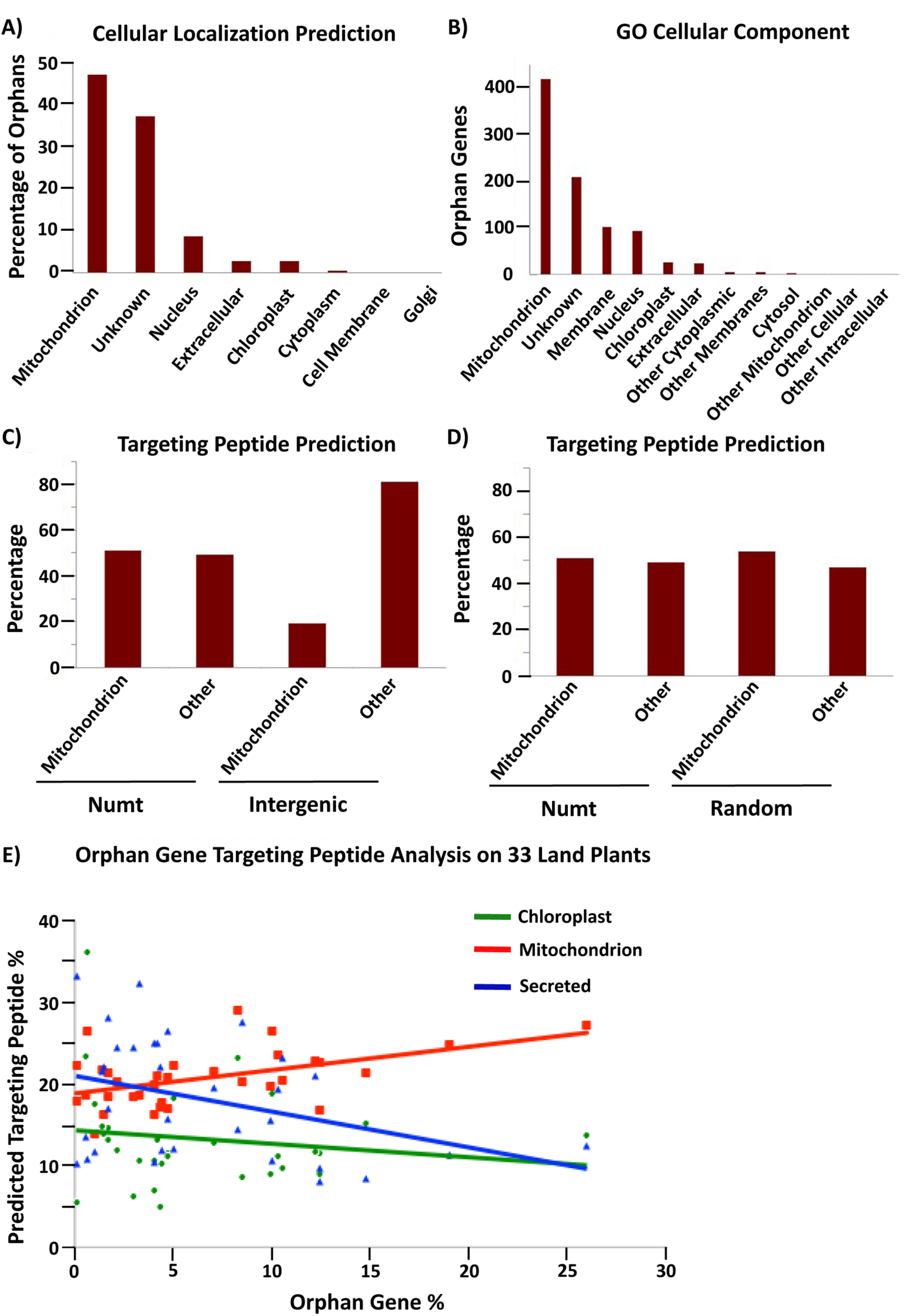
Predicted subcellular localization for *A. thaliana* orphan genes. A) Subcellular localization prediction from TAIR for all *A. thaliana* 1,169 orphan genes. B) Component GO term analysis of *A. thaliana* orphan genes (including 852 orphan genes that have predictions). C) Target peptide analysis of 100 protein sequences generated from *A. thaliana* intergenic and Numt sequences. Numt DNA is more likely to produce a mitochondrial targeting peptide than intergenic DNA (Chi-squared analysis, X-squared = 21.12, df = 1, *P* = 4.312e-06). D) Target peptide analysis on *A. thaliana* Numt DNA and completely randomized DNA show similar proclivities for mitochondrial targeting peptides. E) The percentage of orphan genes with a mitochondrial peptide shows a positive correlation with the percentage of orphan genes in the genome. The correlation coefficient for mitochondria, chloroplast and secreted targeting peptides is 0.488, −0.184, and −0.358 with *P* of 0.004, 0.305, and 0.041 respectively, n = 33.

We next wanted to understand if mitochondrial-originating DNA could preferentially code for a mitochondrial targeting peptide. To do this analysis, a random selection of 100 Numt DNA segments of 174 nucleotides was translated into amino acid sequence. For comparison, a random selection of 100 nuclear intergenic sequences of 174 nt each was translated into amino acid sequence. Each data set was then run through TargetP (Emanualsson et al., 2000) to obtain predicted target peptide sequences. The results revealed that Numt DNA (mitochondrial origin) is more likely to code for a mitochondrial targeting peptide than intergenic DNA (nuclear origin) (Fig. 2C). The same technique was also used for *G. max* and similar results were obtained: Numt DNA from *G. max* was more likely to code for a mitochondrial targeting peptide compared to transposable element (TE) DNA from *G. max* (Fig. S1). The TE DNA was used for comparison as they have long been associated with orphan gene evolution (Bennetzin, 2005). Thus, orphans preferentially target mitochondria, Numt sequences preferentially code for mitochondrial targeting peptides, it is possible that some orphan genes may originate from mitochondrial sequences.

This same analysis was performed on completely randomized DNA sequences (100 sequences of 174 nt). These randomized DNA sequences showed a similar proportion of mitochondrial targeting peptides compared to Numt DNA (Fig. 2D). Thus, mitochondrial-originating DNA seems to have a more random quality than nuclear intergenic DNA as both random DNA sequences and Numt DNA sequences code for a high proportion of mitochondrial targeting peptides.

To further understand how subcellular targeting may relate to orphan gene content, all orphan gene sequences for 33 plant species (from http://www.greenphyl.org/cgi-bin/index.cgi) were run through TargetP to determine their putative targeting peptides, and the percentage of mitochondrion, chloroplast, and secreted targeting peptides were determined. The percentage of orphans with predicted mitochondrial targeting peptides had a positive correlation with the whole genome orphan gene content of each species (Fig. 2E). The other two types of predicted targeting peptides (secreted and chloroplast) had a negative correlation. Therefore, plants with more orphan genes have a higher proportion of orphans with predicted mitochondrial targeting peptides. Since mitochondrial originating DNA can create ORFs that preferentially code for mitochondrial peptides (Fig. 2C and Fig. S1), an increase in orphan genes with mitochondrial targeting peptides may signify an increased mitochondrial gene fostering activity. The correlation between orphan genes and predicted mitochondrial targeting is elucidated further by this analysis of orphan genes of 33 land plant species.

### Orphan gene content is correlated with Numt content in land plant genomes

A collection of Numts for six different land plant species (Numt length data from Ko & Kim et al. 2016) was used to analyze the correlation between mitochondrial DNA transferred to the nucleus and the number of orphan genes present in these species. To do this, the whole Numt length (the total kilobases of every mitochondrial piece of DNA found in the nucleus) was divided by the length of the mitochondrial genome in that species then multiplied by 100 to give a value we termed the Numt turnover %. This tells us how much mitochondrial DNA was transferred in relation to the size of the mitochondrial genome in each species. We then used the total number of orphan genes and the total number of genes in each species to calculate the orphan gene content as a percent of total genes. These two numbers calculated for each of the six species were plotted and analyzed for correlation. Pearson correlation method returned a correlation coefficient of 0.91 with a *P* = 0.01 (Fig. 3A). This provides evidence that there is a positive correlation between the level of mitochondrial DNA transfer (Numt turnover %) and the proportion of orphan genes to total genes in a species (orphan gene %).

**Figure 3.**
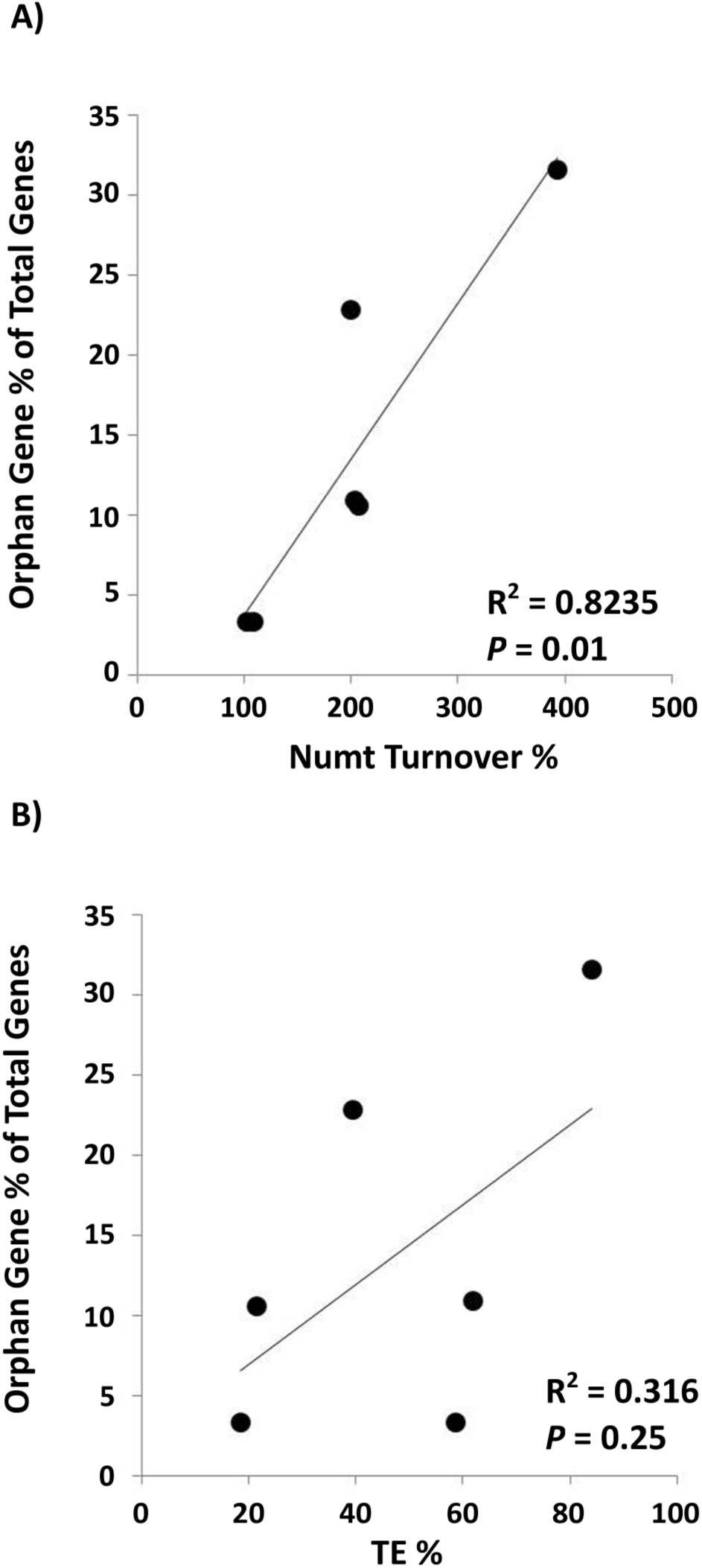
Positive correlation between Numts and orphan genes. A) A significantly positive correlation between Numt turnover % (total Numt length/mitochondrial genome size) and orphan gene percentage (Correlation coefficient = 0.91; *P* = 0.01). B) No significant correlation is found between % of TEs in the genome and orphan gene percentage (Correlation coefficient = 0.56; *P* = 0.25). Species used in this analysis*: A. thaliana*, *G. max*, *Z. mays, O. sativa, S. bicolor*, and *V. vinifera*. TE data was retrieved from Oliver et al. 2013. Orphan gene data for *O. sativa*, *Z. mays* and S. bicolor was obtained from Yao et al., 2017; *A. thaliana* was obtained from Arendsee et al., 2014; *G. max*, and *V. vinifera* were obtained from http://www.greenphyl.org/cgi-bin/index.cgi.

We also computed the correlation between orphan gene content in a genome and the percentage of TEs that make up the genome. A positive correlation was found, however not statistically significant (correlation coefficient = 0.56, *P* = 0.25) (Fig. 3B).

### Orphan gene content is not correlated with Numt content in Cetacean species

As mitochondrial sizes and dynamics are vastly different between mammals and plants (Ko & Kim, 2016; Burger et al., 2003; Barr et al., 2005) (Fig. S2), orphan gene analysis was performed for six cetacean species via a BLAST filtering technique (Donoghue et al. 2011). Although the mammals have larger whole genomes (Fig. 4A), they have much smaller mitochondrial genomes than land plants (Ko & Kim, 2016), and a much smaller orphan gene content than land plants (Fig. 4B). No correlation was detected between orphan gene content with Numt turnover in mammal genomes (Fig. 4C). This further provides evidence that the mitochondrial genome may play a role in orphan gene development in land plants, while that role has likely been minimized or lost in animals due to their small mitochondrial genome size.

**Figure 4.**
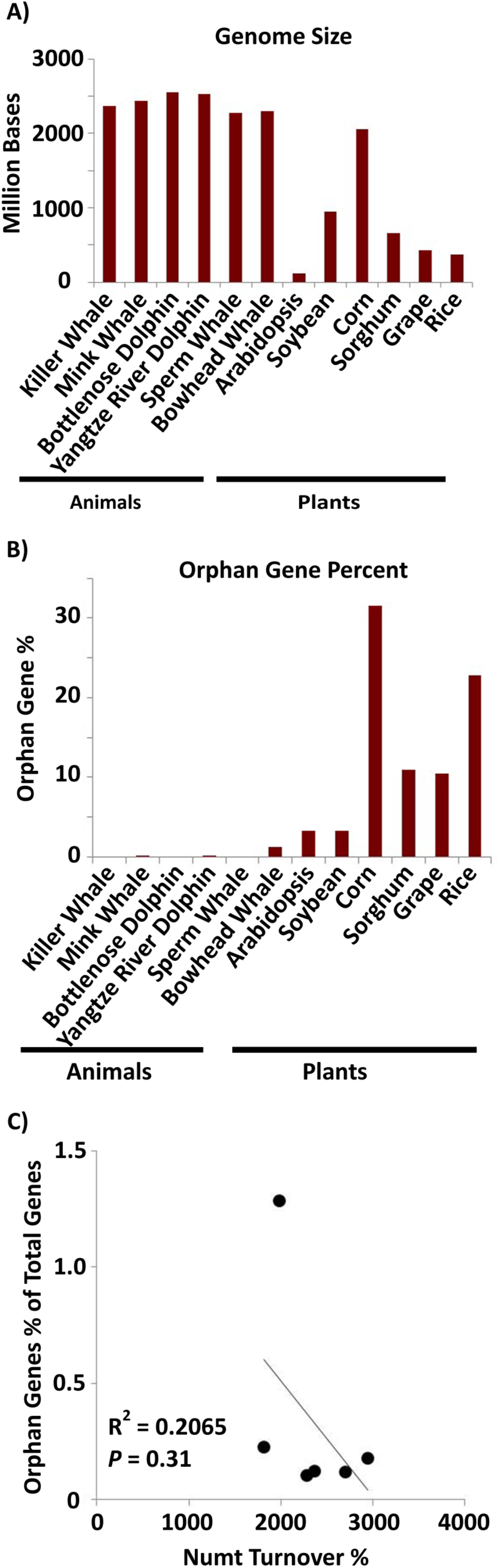
Orphan gene content in six aquatic mammals. A) Whole genome sizes of the six mammalian species and the six plant species (data from NCBI). B) Orphan gene content (as a percentage of total genes) for aquatic mammals and land plants. C) The correlation of orphan gene content as percentage of total genes and Numt turnover %. Correlation coefficient = −0.48; *P* = 0.31. The amount of orphan genes in mammal species is not significantly correlated with their Numt content.

### *A. thaliana* orphan genes in both mitochondrial genome and nuclear genome

Of the 30 predicted mitochondrial orphan genes in *A. thaliana*, 28 of them have at least some homology to a nuclear chromosome (Table 2). Of these 28, 22 orphans have a 100% query cover to a nuclear chromosome which means the entire sequence can be detected in both mitochondrial genome and nuclear genome, thus it is possible that the sequence could have been transferred from the mitochondrial genome to the nuclear genome. Most of these transfers are seen in chromosome two—which may be a result of the recent, large transfer event where nearly the entire mitochondrial genome was transferred to the pericentromeric region of the chromosome (Stupar et al., 2001). Furthermore, 29 of the 30 have hits to mitochondrial genome in another species. This implies that the sequences originated in the mitochondrial genome, and were recently transferred to the nuclear genome of *A. thaliana*. Only one orphan had sequence homology to a nuclear genome in a species other than *A. thaliana*. This reinforced the idea that these sequences originated in the mitochondrial genome.

**Table 2.**
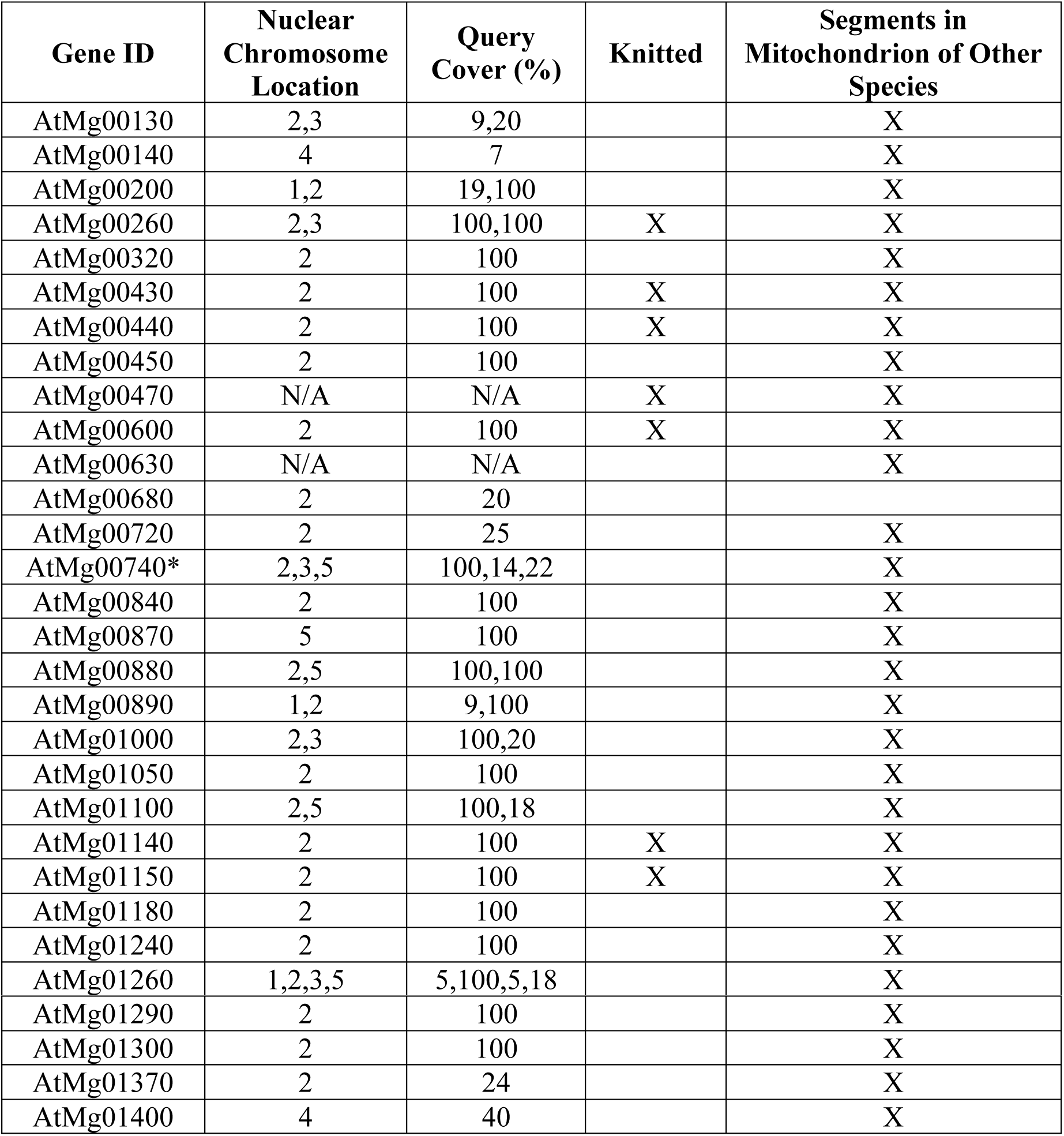
Genome homology of orphans of mitochondrial genome in A. thaliana. Nearly all predicted orphan genes in the mitochondria have either complete or partial sequence transfer to the nuclear genome of Arabidopsis. *AtMg00740 is the only gene that has sequence homology to a nuclear chromosome of a different species (*A. alpina*). All genes except AtMg00680 have some sort of sequence homology to another species mitochondrial genome, suggesting the sequence originated in the mitochondrial genome and has been recently transferred to the nuclear genome of *A. thaliana*. Orphans were marked as knitted if two different segments of the query was found in the same mitochondrial genome of a particular species (other than *A. thaliana*).

| Gene ID | Nuclear Chromosome Location | Query Cover (%) | Knitted | Segments in Mitochondrion of Other Species |
| --- | --- | --- | --- | --- |
| AtMg00130 | 2,3 | 9,20 |  | X |
| AtMg00140 | 4 | 7 |  | X |
| AtMg00200 | 1,2 | 19,100 |  | X |
| AtMg00260 | 2,3 | 100,100 | X | X |
| AtMg00320 | 2 | 100 |  | X |
| AtMg00430 | 2 | 100 | X | X |
| AtMg00440 | 2 | 100 | X | X |
| AtMg00450 | 2 | 100 |  | X |
| AtMg00470 | N/A | N/A | X | X |
| AtMg00600 | 2 | 100 | X | X |
| AtMg00630 | N/A | N/A |  | X |
| AtMg00680 | 2 | 20 |  |  |
| AtMg00720 | 2 | 25 |  | X |
| AtMg00740* | 2,3,5 | 100,14,22 |  | X |
| AtMg00840 | 2 | 100 |  | X |
| AtMg00870 | 5 | 100 |  | X |
| AtMg00880 | 2,5 | 100,100 |  | X |
| AtMg00890 | 1,2 | 9,100 |  | X |
| AtMg01000 | 2,3 | 100,20 |  | X |
| AtMg01050 | 2 | 100 |  | X |
| AtMg01100 | 2,5 | 100,18 |  | X |
| AtMg01140 | 2 | 100 | X | X |
| AtMg01150 | 2 | 100 | X | X |
| AtMg01180 | 2 | 100 |  | X |
| AtMg01240 | 2 | 100 |  | X |
| AtMg01260 | 1,2,3,5 | 5,100,5,18 |  | X |
| AtMg01290 | 2 | 100 |  | X |
| AtMg01300 | 2 | 100 |  | X |
| AtMg01370 | 2 | 24 |  | X |
| AtMg01400 | 4 | 40 |  | X |

### Mitochondrial fostering of At2g07667 (*KNIT*), a young *A. thaliana* gene

To better understand how orphan sequences may evolve in the mitochondrial genome, we analyzed a predicted nuclear orphan gene that has sequence homology to the mitochondrial genome of *A. thaliana*. Gene At2g07667, termed here as Knitted In Mitochondria (*KNIT*), is a protein-coding gene with two proposed gene models. There are six exons, and five introns in each of the models (Fig. 5A). The protein consists of 218 amino acids and its full-length genomic sequence covers 1,746 base pairs (<u>arabidopsis.org</u>). Alignment analysis carried out with the NCBI BLASTn server reveals the nucleotide sequence of this gene is also found in the mitochondrial genome of *A. thaliana* (identity and query cover of 100%, Fig. 5A). A BLASTp search revealed that the KNIT protein only has one potential homolog in *Brassica napus* (sequence ID: CDY45527.1), a protein made up of 171 amino acids with a query cover of 76% and an identity of 77%. tBLASTn analysis on the peptide sequence of the *B. napus* homolog shows that the protein maps back to the mitochondrial genome of *B. napus* covering 3 different ranges, suggesting at least two introns. This hit (BLASTp) to *B. napus* indicates that *KNIT* may not be a “pure” species-specific orphan gene, but is indeed a very young gene.

**Figure 5.**
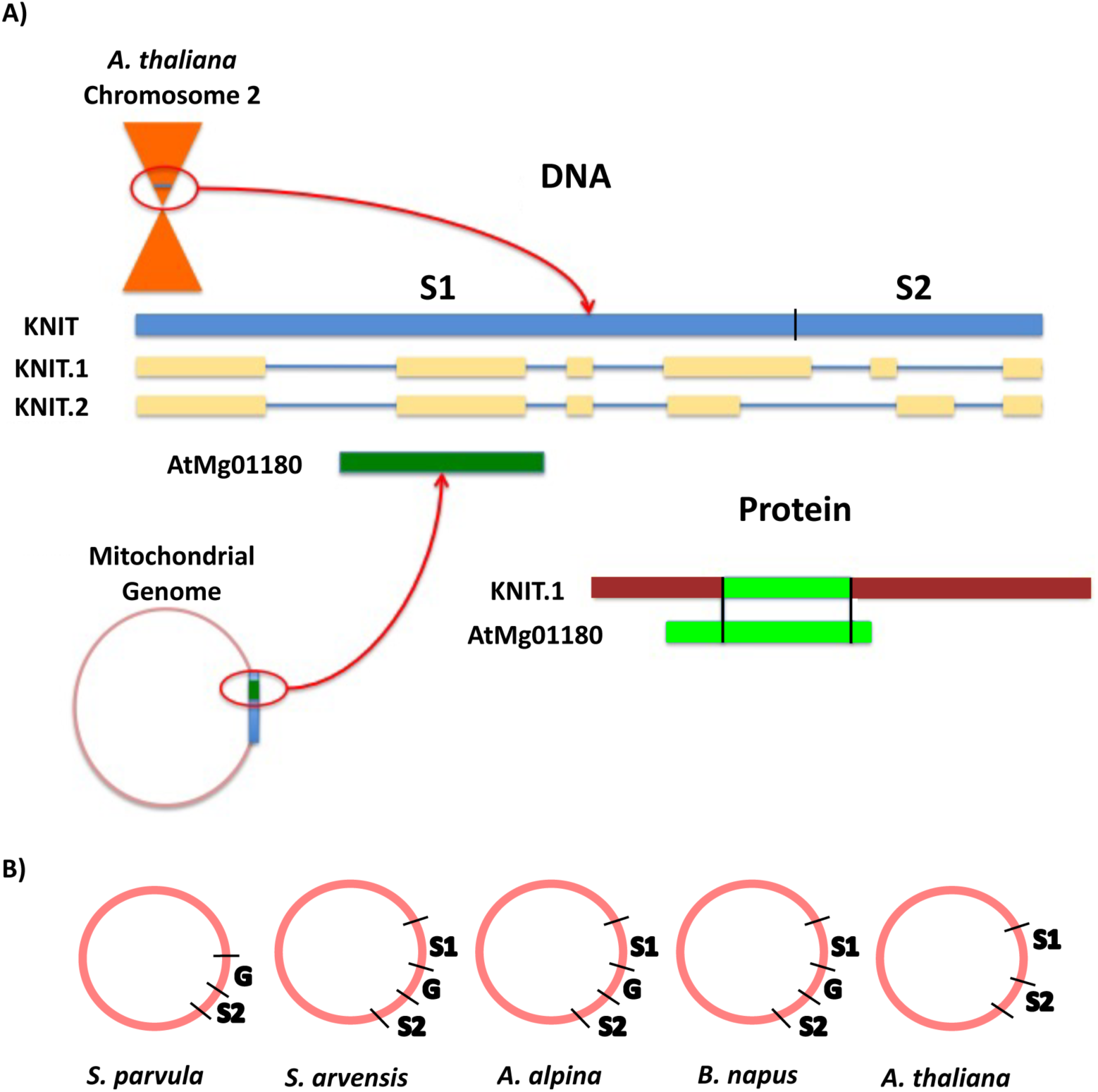
KNIT sequence origin and location. A) Model of KNIT loci in both chromosome 2 and the mitochondrial genome. The model indicates that the full genomic sequence of KNIT exists in both chromosome two near the centromere and in the mitochondrial genome of *A. thaliana*, the predicted protein sequence differ. B) The gap sequence that separates genic sequences is not present in *A. thaliana*. G= gap sequence; S1= first segment of *KNIT* gene; S2= second segment of *KNIT* gene.

The only other protein sequence hit for the query KNIT belongs to the *A. thaliana* mitochondrial genome. However, even though the entire nucleotide sequence of *KNIT* is found in the mitochondrial genome, the gene annotated in this region is not identical to *KNIT*. In the mitochondrial genome, the gene locus ID AtMg01180 (Table 2), codes for a protein of 111 amino acids covering a portion of the first intron of KNIT, the entire second exon, and a portion of the second intron—covering amino acids 64-120 of KNIT protein (Fig. 5A). Thus, the same nucleotide sequence is expressed in different ways between the mitochondrial and the nuclear genomes.

Alignment analysis (BLASTn) of KNIT showed hits throughout different mitochondrial genomes: *Schrenkiella parvula, Arabis alpina, Brassica napus, Sinapis arvensis*, and of course *A. thaliana* (Table 3). This suggests that the gene sequence originated in the mitochondrial genome. This sequence was possibly transferred to the *A. thaliana* chromosome two via a recent transfer event (Stupar et al., 2001). The first 1,290 nucleotides (Segment 1) of KNIT can be found in all above species except *S. parvula*, and the second 475 nucleotides (Segment 2) is found in all 5 species. However, these two gene segments are separated by a 652 nucleotide segment (Gap) in *A. alpine, B. napus*, and *S. arvensis*. This Gap sequence is not present in *A. thaliana*, but is present in *S. parvula* as well (Fig. 5B). Along with KNIT, seven of the mitochondrial orphans show clear knitting patterns—multiple segments of the query were found in multiple locations in the mitochondrial genome (Table 2).

**Table 3.**
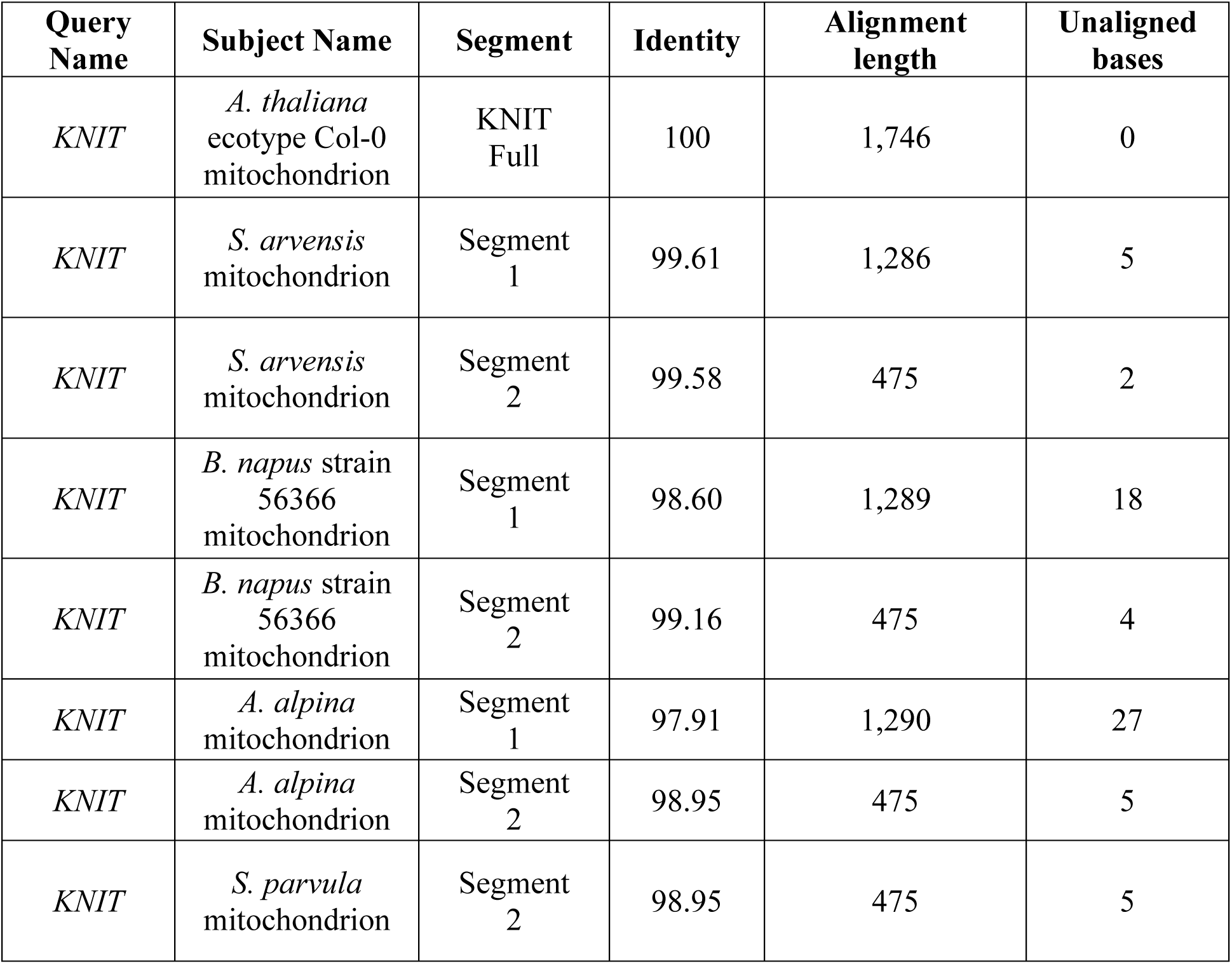
Alignment analysis for KNIT (BLASTn hit). One hit was reported in *A. thaliana* (outside of the chromosome 2 hit) with a 100% match spanning all 1,746 nucleotides. For *S. arvensis*, *B. napus*, and *A. alpina*, two individual hits (one around 1,290 nt and the other 475 nt) to the mitochondrial genome were reported. *S. parvula* had one hit (475 nt) reported. A small amount of unaligned bases for all hits underlie a low mutation rate of the mitochondrial genomes.

| Query Name | Subject Name | Segment | Identity | Alignment length | Unaligned bases |
| --- | --- | --- | --- | --- | --- |
| *KNIT* | *A. thaliana* ecotype Col-0 mitochondrion | KNIT Full | 100 | 1,746 | 0 |
| *KNIT* | *S. arvensis* mitochondrion | Segment 1 | 99.61 | 1,286 | 5 |
| *KNIT* | *S. arvensis* mitochondrion | Segment 2 | 99.58 | 475 | 2 |
| *KNIT* | *B. napus* strain 56366 mitochondrion | Segment 1 | 98.60 | 1,289 | 18 |
| *KNIT* | *B. napus* strain 56366 mitochondrion | Segment 2 | 99.16 | 475 | 4 |
| *KNIT* | *A. alpina* mitochondrion | Segment 1 | 97.91 | 1,290 | 27 |
| *KNIT* | *A. alpina* mitochondrion | Segment 2 | 98.95 | 475 | 5 |
| *KNIT* | *S. parvula* mitochondrion | Segment 2 | 98.95 | 475 | 5 |

To explain the possible impact of the mitochondrial genome on orphan gene evolution, we propose a model with two routes whereby orphan open reading frames are either: 1) generated in the mitochondrial genome and transferred to the nuclear genome as a Numt or 2) mitochondrial DNA is transferred to the nuclear genome and further diversification (*i.e.*, cutting by TEs) generates novel open reading frames (Fig. 6). Based on the large percentage of orphan genes in plant mitochondria, and evidence that these open reading frames can transfer among genomes, it is likely that Route 1 may play a role in the evolution of novel genic content in the nuclear genome.

**Figure 6.**
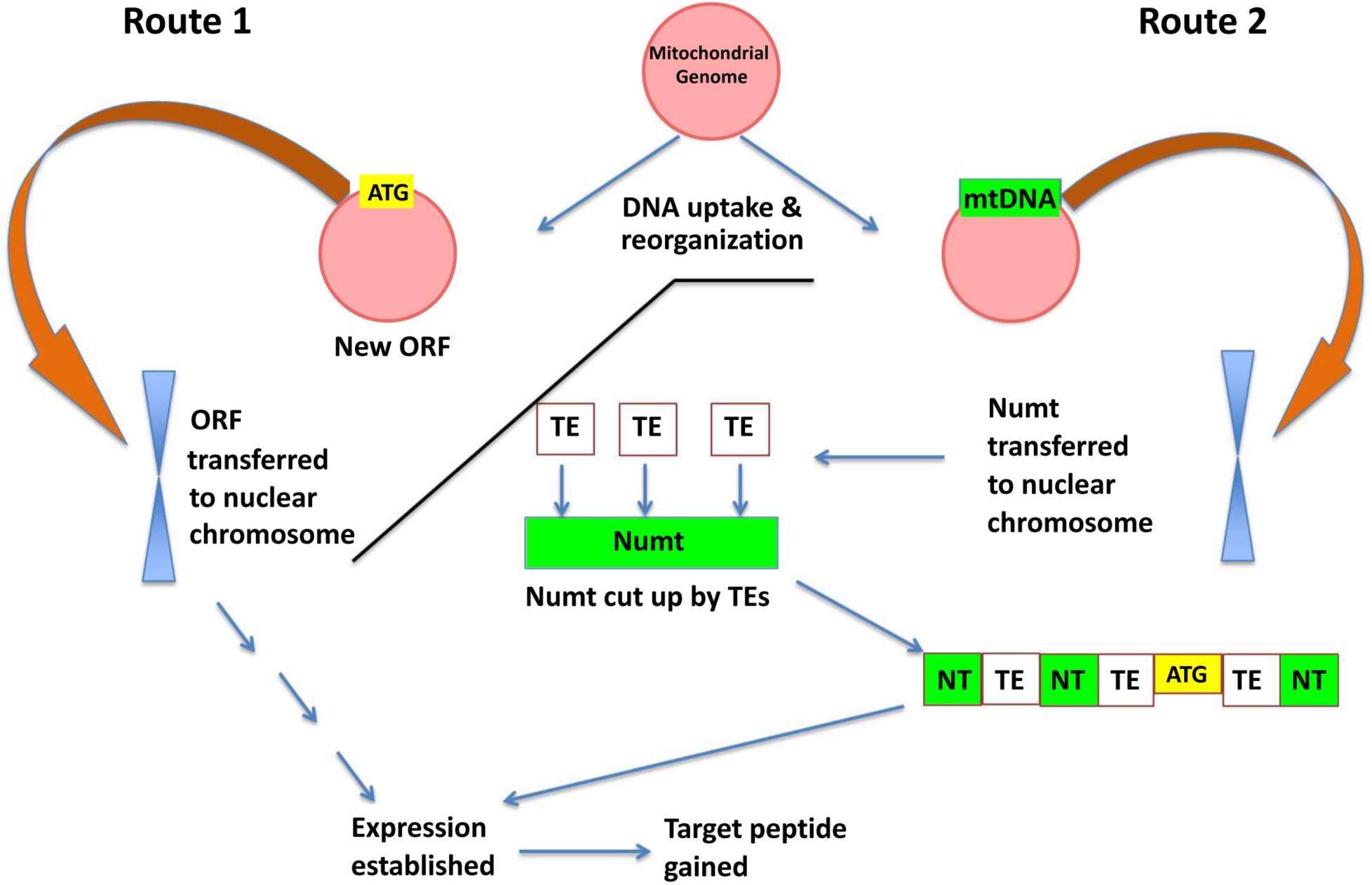
A model for orphan gene evolution through the mitochondrial genome. Novel sequences are created due to the high rearrangement rate in the mitochondrial genome, and then inserted into a nuclear chromosome. The transferred DNA may already contain gene coding information like *KNIT* (left side of model: Route 1) or may obtain an open reading frame via other genome mechanisms such as transposon transposition (right side: Route 2).

## Discussion

We analyzed orphan gene content in three well annotated plant mitochondrial genomes and found the mitochondrial genome contains a higher proportion of orphan genes compared to the chloroplast genome and whole genome (Fig. 1). This is likely because of the high recombination rates and sequence diversity of the plant mitochondrial genome (Christensen, 2013). As large mitochondrial sequences are often transferred to the nucleus (Ko et al., 2015; Lopez et al., 1996; Michalovova et al., 2013; Richly & Leister, 2004; Noutsos et al., 2005; Bensasson et al., 2001; Stupar et al., 2001), looking into the relationship between Numts and orphan genes may prove fruitful in understanding *de novo* gene evolution.

As the original transfer of mitochondrial sequences to the nuclear genome led to a mass organelle gene exodus that required the transferred genes to obtain targeting back to the organelle (Berg & Kurland, 2000), we examined whether current orphan genes in *A. thaliana* preferentially target mitochondria. Indeed, we found that a large proportion of orphan genes are predicted to target mitochondria (Fig. 2). We next determined if mitochondrial originating DNA (Numts) preferentially code for mitochondrial targeting peptides compared to intergenic DNA. We found that mitochondrial originating DNA does preferentially code for a mitochondrial targeting peptide (Fig. 2). This further leads us to believe that it is possible for orphan genes in *A. thaliana* to arise from mitochondrial originating DNA. Mitochondrial originating DNA and completely randomized DNA sequences each preferentially code for mitochondrial targeting peptides, suggesting that Numts have a random quality, possibly allowing for the creation of novel genes with no known functional motifs, a trait common to orphan genes (Arendsee et al., 2014) (Fig. 2).

When looking at the genomes as a whole, mammal genomes are much larger, while their mitochondrial genomes are much smaller than plant mitochondrial genomes (Fig. 4A). We can also see that plant mitochondrial genomes are much more dynamic in size between the organisms whereas mammal mitochondrial genomes are nearly identical in size (Fig. S2). Plants also have more Numts on average than animals (Bensasson et al., 2001), and they appear to be more dynamic in size as well (Ko & Kim, 2016). Therefore, mammal mitochondrial genomes, having lost size and dynamic nature, likely do not play as much of an active role in orphan gene evolution, hence the smaller amount of orphan genes and lack of correlation with Numts compared to plant mitochondrial genomes (Fig. 3&4).

To determine if full orphan gene sequences can transfer from the mitochondrial genome to the nuclear genome, sequence alignment analysis was performed for all 30 predicted mitochondrial orphan genes in *A. thaliana*. The vast majority of these orphan sequences were also found in the nuclear chromosomes of *A. thaliana*. Hits to species other than *A. thaliana* were limited to the mitochondrial genomes (Table 2). This indicates these orphans were formed in the mitochondrial genome and their sequences were recently transferred to the nuclear genome.

In order to better understand how the mitochondrial genome can create orphan genes and subsequently transfer them to the nuclear genome, an orphan gene with a sequence in both the mitochondrial genome and chromosome two was chosen for sequence analysis. This analysis appears to show the evolution of *KNIT* as a result of reshuffling in the mitochondrial genome of *A. thaliana* (Fig. 5). The low number of SNPs (single nucleotide polymorphism) between the partial *KNIT* sequences in the mitochondria of *A. thaliana* and other species (Table 3) further demonstrates that the creative power of the mitochondrial genome is in its rearrangement ability and not its mutation rate. Also, like *KNIT*, several of the mitochondrial orphan genes have evidence of being knitted in the mitochondrial genome due to genome rearrangements (Table 2). Based on this work and the current knowledge behind plant mitochondrial genome dynamics, we propose a model (Fig. 6) whereby novel sequences, sometimes containing orphan genes, are generated in the plant mitochondrial genomes based on uptake of DNA from the nucleus/chloroplast/horizontal gene transfer and subsequent reshuffling of the organelle genome. When large Numts are transferred to the nuclear genome, the novel sequences are transferred as well and are then subject to evolutionary mechanisms in the nuclear genome (higher mutation rate/enhanced epigenetic control/TEs/etc.). Mitochondrial fostering may play an integral role in the evolution of *de novo* orphan genes.

## Methods

### *A. thaliana, G. max and Z. mays* orphan gene analysis

Orphan gene number for the whole genome was gathered from <u>greenphyl.org</u> for *G. max*, from Yao et al., 2017 for *Z. mays*, and from Arendsee et al., 2014 for *A. thaliana*. Orphans were determined for the mitochondrial genome and the chloroplast genome using OrfanFinder (Ekstrom & Yin, 2016).

### *A. thaliana* targeting peptide and Numt analysis

All analysis was performed with the *A. thaliana* genome from NCBI. Accession numbers: chromosome 1, NC_003070.9; chromosome 2, NC_003071.7; chromosome 3, NC_003074.8; chromosome 4, NC_003075.7; chromosome 5, NC_003076.8; and mitochondrial genome, NC_037304.1.

Using NCBI’s BLASTn, a list of Numts was generated by aligning the mitochondrial genome of Arabidopsis against the nuclear genome of Arabidopsis using parameters from Ko & Kim, 2016. Numt fasta files were obtained using the getfasta command-line tool of the Bedtools suit, and Numts within 10,000 bps were merged together via mergefasta from Bedtools (Quinian & Hall, 2010). A similar procedure was used for *G. max* Numt determination. From this list we sampled 100 random sequences of 174 nucleotides long (translated into a 60-aa-long protein, a typical length for young proteins; cutting into 174 nucleotide segments was performed at <u>bioinformatics.org</u>). We randomly sampled 100 non-coding nuclear DNA sequences of 174 nt long. To do this, intergenic regions were collected (regions of the genome which do not code for any genes) from Araport (<u>araport.org</u>). Both lists were run through the TargetP server (www.cbs.dtu.dk/services/TargetP/) to determine their putative targeting peptides. For each sequence, we received a letter that represents the presence of the potential targeting sequence (M = mitochondria, C = chloroplast, S = secreted, O = other). Data was organized in Excel. Statistical analysis was performed with R. For *G. max*, we compared Numt DNA with random TE sequences. TE sequence data was obtained from Soybase (https://www.soybase.org/soytedb/).

### Targeting peptide analysis on orphan genes from 33 plant species

Orphan gene protein FASTA sequences for 33 plant species were downloaded from <u>greenphyl.org</u> and TargetP was used to predict targeting peptides. Analysis of TargetP data was performed with in-house python code.

### Correlation of Numts and orphan genes in six plant species

Orphan gene data for *O. sativa*, and *S. bicolor* was obtained from Yao et al., 2017, from http://www.greenphyl.org/cgi-bin/index.cgi for *V. vinifera*. Numt length and mitochondrial genome size were obtained from Ko & Kim, 2016.

### Numt and Orphan gene analysis in Cetaceans

In order to determine orphan gene content in the six Cetacean species, we used a two-step BLAST filtering method. First, we used local diamond BLASTp (Buchfink et al., 2015), a BLAST algorithm that can perform up to 20,000 times as fast as NCBI BLASTp, to align our query proteome to the other five Cetacean proteomes (E-value = 0.001). The unaligned query sequence IDs were reported back to us. This gave us our initial set of prospective orphan genes.

Next, we ran this list of orphans through NCBI tBLASTn against the refseq_rna database (E-value = 0.01). We excluded searching to mRNA from our query species. After this step, our unaligned query IDs became our final orphan gene list.

Proteome sequences were gathered from Uniprot (https://www.uniprot.org/uniprot/) for all species except the minke whale (NCBI) and the Bowhead whale (http://www.bowhead-whale.org/). Diamond BLASTp was carried out locally under default search parameters (E-value cutoff of 0.001) and databases were created manually. Query files for tBLASTn were run on the NCBI server with an E-value cutoff of 0.01. Whole genome size for plant species was found at NCBI.

### Mitochondrial orphan gene analysis

Full genomic DNA sequences for the orphan genes were downloaded from TAIR. Alignment analysis was carried out on the NCBI BLASTn webserver. If two different segments of a query sequence were found in two different regions of mitochondrial genome of another species (other than *A. thaliana*), the query sequence was considered knitted.

### KNIT analysis

KNIT full length genomic DNA sequence was downloaded from TAIR (<u>arabidopsis.org</u>). Alignment analysis was carried out on the NCBI BLAST webserver.

## Acknowledgements

We thank Lei Wang and Rezwan Tanvir for helpful discussions. This study is supported by funding from Mississippi State University to LL.

## Author contributions statement

SO conceived the study, designed and performed the experiments, analyzed the data, and wrote the paper. LL conceived the study, designed the experiments, and wrote the paper.

