## Supplementary Figures for "Mitochondrial fostering: the mitochondrial genome may play a role in plant orphan gene evolution"

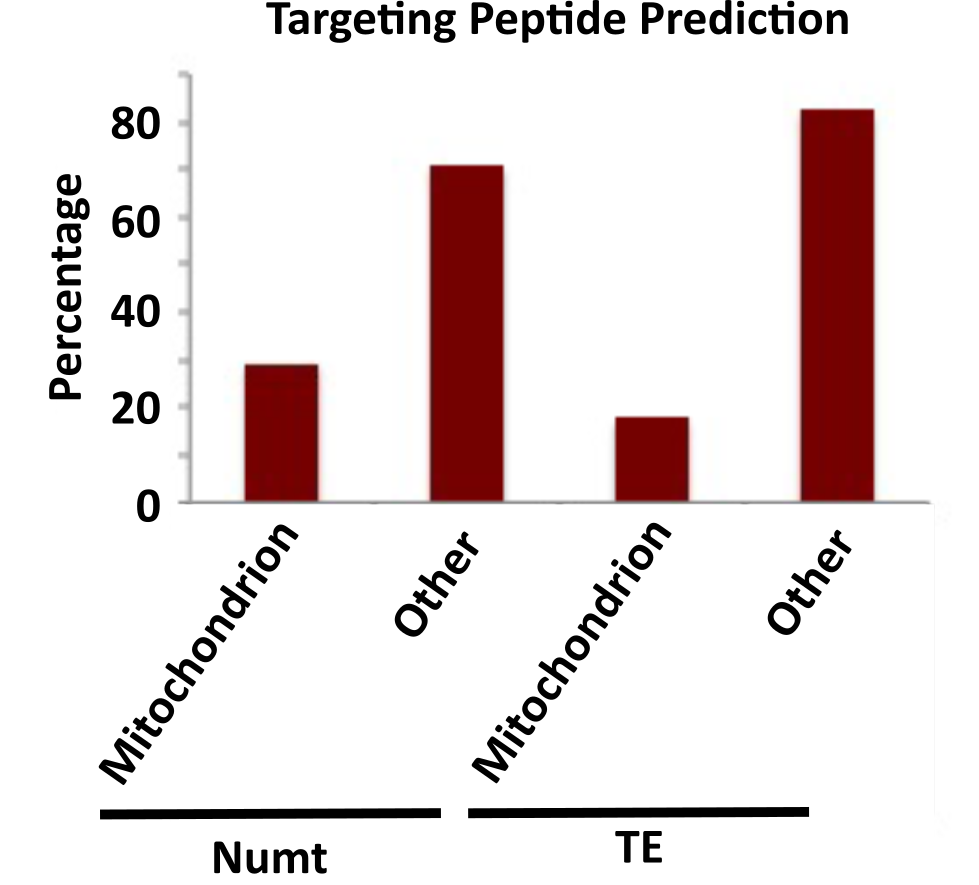

**Figure S1.** One hundred sixty-four sequences of 174 nt were randomly sampled for *G. max* Numt DNA and transposable element DNA, and subsequently run through TargetP. As similar as in Arabidopsis, Numt DNA in *G. max* is more likely to code for a mitochondrial targeting peptide as compared to transposable element DNA (Chi-squared *P* = 0.0001).

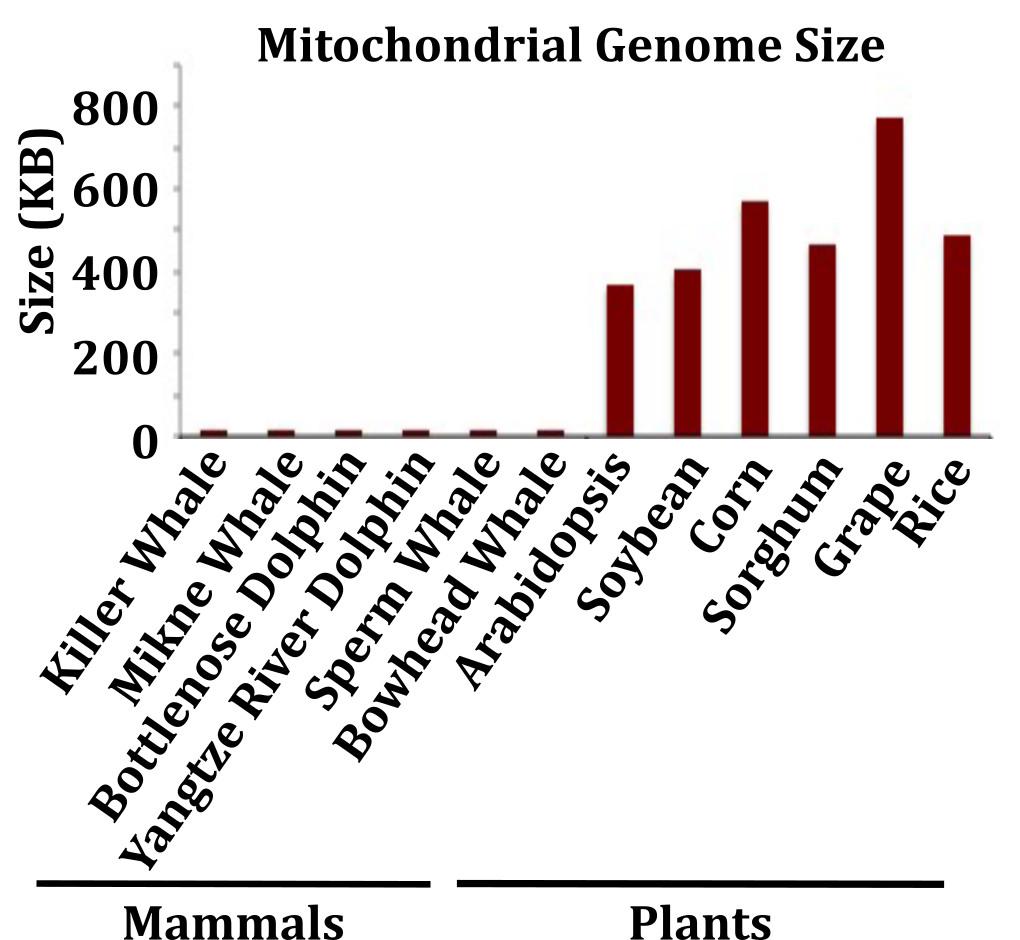

**Figure S2.** **Mitochondrial genome size for six mammal species and six plant species.** Mitochondrial genomes are much larger in plant species compared to the mammal species used in this study.
